# Selective regulation of corticostriatal synapses by astrocytic phagocytosis

**DOI:** 10.1101/2024.03.26.586775

**Authors:** Ji-young Kim, Hyeyeon Kim, Won-Suk Chung, Hyungju Park

## Abstract

In the adult brain, neural circuit homeostasis depends on the constant turnover of synapses via astrocytic phagocytosis mechanisms. However, it remains unclear whether this process occurs in a circuit-specific manner. Here, we reveal that astrocytes target and reorganize excitatory synapses in the striatum. Using model mice lacking astrocytic phagocytosis receptors in the dorsal striatum, we found that astrocytes constantly remove corticostriatal synapses rather than thalamostriatal synapses. This preferential elimination suggests that astrocytes play a selective role in modulating corticostriatal plasticity and functions via phagocytosis mechanisms. Supporting this notion, corticostriatal long-term potentiation (LTP) and the early phase of motor sequence learning are dependent on astrocytic phagocytic receptors. Together, our findings demonstrate that astrocytes contribute to the connectivity and plasticity of the striatal circuit by preferentially engulfing a specific subset of excitatory synapses within brain regions innervated by multiple excitatory sources.

## Introduction

Proper synaptic connectivity is vital for normal brain functioning. A growing body of evidence suggests that glial phagocytosis plays a crucial role in maintaining appropriate synaptic connections^1–3^. In addition to its role in refining neural circuits during development^4–7^, recent studies have highlighted the direct involvement of glial synapse reorganization in mature neural circuits^8,9^. This glia-mediated synapse reorganization in the adult brain is essential for normal cognitive functions, including learning and memory^8,9^.

Astrocytes, a primary type of glial cell, actively contribute to synapse reorganization by phagocytosing target synapses via MEGF10 (multiple epidermal growth factor-like domains protein 10) and MERTK (Mer proto-oncogene tyrosine kinase) phagocytic receptors^7,9^. Apart from their involvement in the maturation of developing neural circuits^7^, MEGF10/MERTK-dependent astrocytic phagocytosis also play an important role in maintaining hippocampal neural networks, which are required for memory formation in the adult brain^9^. However, it remains unclear whether astrocytic phagocytosis-dependent synapse reorganization occurs uniformly across all mature neural circuits or exhibits circuit specificity.

The striatum, an important brain region governing procedural and motor skill learning, receives multiple innervations from various brain areas, such as the cortex and thalamus^10,11^. The experience-dependent plasticity of cortico- and thalamostriatal pathways plays a crucial role in motor learning^12–15^. While the thalamostriatal pathway is involved in execution, the corticostriatal pathway is primarily involved in motor skill learning^15^. The distinct roles of these pathways raise the possibility that they are differentially modulated, with astrocytes potentially playing a role in achieving proper functional integration. Because of the region-specific molecular heterogeneity between hippocampal and striatal astrocytes^16^, striatal synapses may be modulated via different astrocytic synapse reorganization mechanisms than hippocampal synapses. However, comparable expression levels of MEGF10 and MERTK were found in both hippocampal and striatal astrocytes (http://astrocyternaseq.org/adultastro)^16^, suggesting that striatal astrocytes and hippocampal astrocytes are regulated by the same synapse reorganization mechanisms. Since the role of either of these two phagocytic receptors in eliminating unnecessary striatal synapses has never been tested, we decided to utilize a *Megf10/Mertk* double-knockout strategy to examine the exact contribution of astrocytic phagocytosis to striatal functions.

In this study, we showed that astrocytic *Megf10/Mertk*-dependent phagocytosis targets and reorganizes striatal excitatory synapses. Moreover, astrocytic synapse refinement was more frequent at corticostriatal synapses than at thalamostriatal synapses. These results suggest that astrocytic phagocytosis reorganizes excitatory synapses in a circuit- specific manner.

## Results

### Astrocytic MEGF10/MERTK signaling is required for the elimination of excitatory synapses in the adult striatum

To test whether astrocytic phagocytosis is involved in synapse turnover in the adult striatum, we first examined whether striatal excitatory or inhibitory synaptic transmission is influenced by impaired astrocytic phagocytosis. To efficiently inhibit astrocyte-mediated phagocytosis, the astrocyte phagocytosis receptors *Megf10* and *Mertk*^7,9^ were selectively deleted in striatal astrocytes using double-loxP-floxed *Megf10* and *Mertk* (*Megf10*^fl/fl^; *Mertk*^fl/fl^) mouse model (Depletion of Astrocytic Phagocytosis in the Striatum; DAPS). The DAPS model was established by injecting an adeno- associated virus (AAV) encoding GFAP (0.7) promoter-driven Cre (AAV-GFAP_0.7_- EGFP-T2A-iCre) into the dorsal striatum of *Megf10*^fl/fl^; *Mertk*^fl/fl^ mice. For controls, the same mice were injected with control AAV (AAV-GFAP_0.7_-EGFP) (Fig. 1a). Virus expression was confirmed to be astrocyte-specific in both control and DAPS groups, with a significant reduction in MEGF10 and MERTK expression observed in the DAPS group (Supplementary Fig. 1).

**Figure 1.**
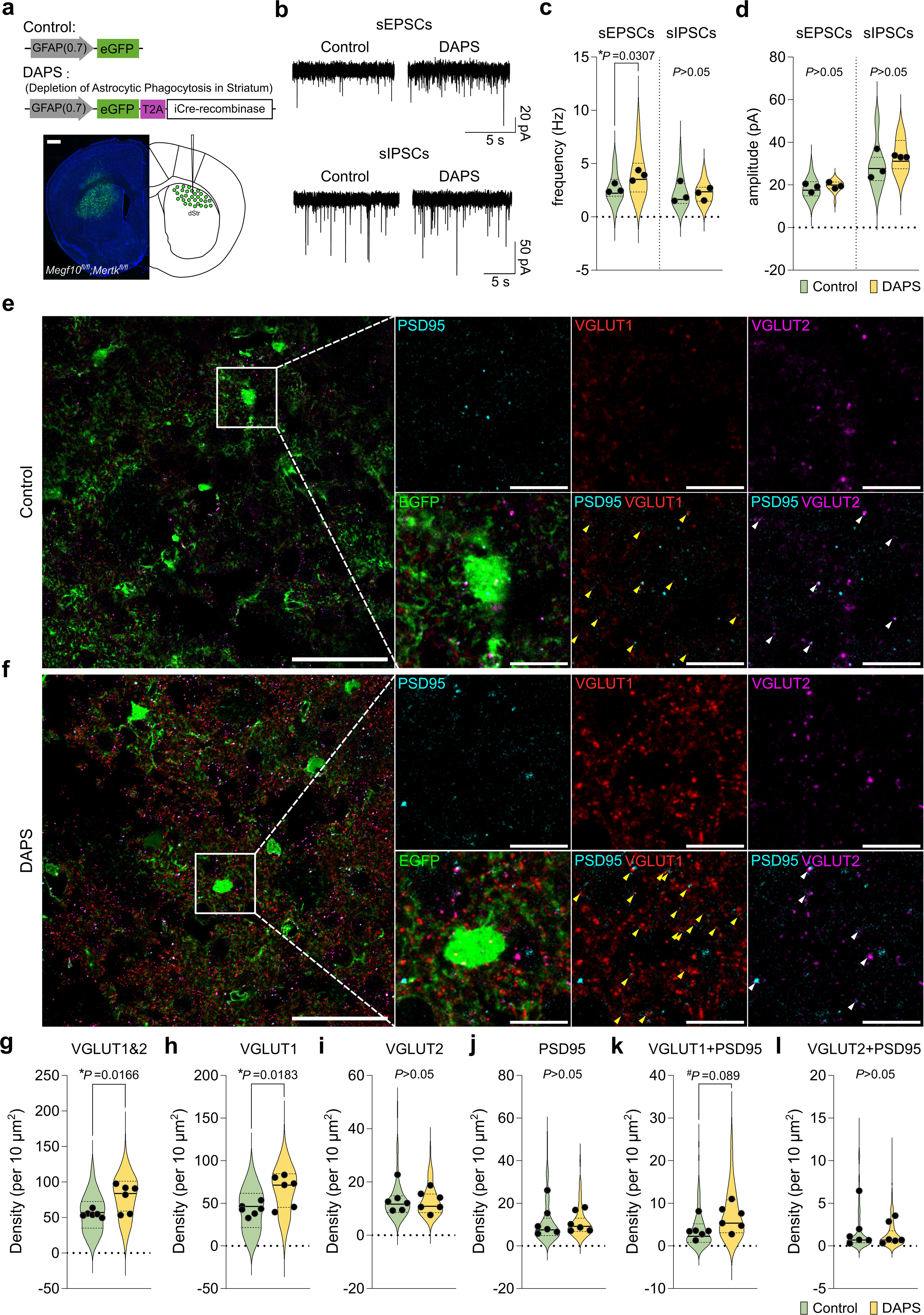
Selective regulation of excitatory synapses by astrocytic MEGF10/MERTK in the adult dorsal striatum. **a.** Schematic diagram showing the DAPS mouse model. **b.** Representative traces showing spontaneous EPSCs or IPSCs recorded from MSNs from the control and DAPS groups. **c-d.** Summary of frequencies (**c**) or amplitudes (**d**) of sEPSCs and sIPSCs. Control, n= 15 cells from 3 mice; DAPS, n= 16 cells from 3 mice for sEPSC experiments. Control, n= 21 cells from 3 mice; DAPS, n= 17 cells from 3 mice for sIPSC analysis. **e-f.** Representative confocal images depicting excitatory synapse markers in the dorsal striatum of control (**e**) and DAPS (**f**) mice. AAV-infected astrocytes (EGFP), VGLUT1 , VGLUT2 , and PSD-95 are shown. Left, representative single-plane images. Scale bars = 50 μm; right, expanded views of the indicated white box. Scale bars = 10 μm. Yellow arrows indicate colocalization of VGLUT1 and PSD-95. The white arrows indicate colocalization of VGLUT2 and PSD95. **g-l.** Summary of the density of VGLUT1&2 (**g**)-, VGLUT1 (**h**)-, VGLUT2 (**i**)-, PSD-95 (**j**)-positive particles and VGLUT1 and PSD95 (**k**)- and VGLUT2 and PSD95 (**l**)-double-positive particles. Control, n= 130 images from 6 mice; DAPS, n= 131 images from 6 mice. Unpaired student’s t-test. Mean ± s.e.m.

Both spontaneous excitatory and inhibitory postsynaptic currents (sEPSCs and sIPSCs) were recorded from medium spiny neurons (MSNs) in the dorsal striatum to detect changes in excitatory and inhibitory network activity, respectively. While no alterations were observed in the membrane properties of MSNs in DAPS mice (Table 1), we found that the average frequency of sEPSCs in DAPS striatal slices was significantly greater than that in control slices (Fig. 1b, c). However, the amplitudes of sEPSCs and both the frequencies and amplitudes of sIPSCs remained unchanged in the DAPS slices (Fig. 1b- d). These results suggest that astrocytic regulation of excitatory synaptic transmission occurs through phagocytosis mechanisms.

Immunohistochemical analysis of striatal synapses revealed alterations in excitatory synapses in the DAPS mouse brain slices. When the numbers of excitatory or inhibitory synapses was quantified in the dorsal striatum of control or DAPS mice, we found that while PSD-95 density remained unchanged (Fig. 1e, f, j), the number of excitatory presynaptic markers (estimated by the numbers of VGLUT1- and VGLUT2-density) was significantly increased in the DAPS striatum (Fig. 1g, Supplementary Fig. 2a). However, the control and DAPS striatum exhibited no differences in the numbers of both VGAT- and gephyrin-positive puncta (Supplementary Fig. 2b-e), indicating that astrocytic phagocytosis has no impact on the number of inhibitory synapses. These biased effects of astrocytic phagocytosis on excitatory synapses are consistent with previous findings that hippocampal excitatory synapses are more sensitive to astrocytic phagocytosis mechanisms than are inhibitory synapses^9^.

### Preferential regulation of corticostriatal synapse numbers by astrocytic phagocytosis

The dorsal striatum receives afferent glutamatergic inputs from diverse sources, including the cortex and thalamus^10,11^. Because both cortico- and thalamostriatal excitatory synapses are important for regulating procedural and motor learning and memory, we next investigated whether striatal astrocytes phagocytose these two types of excitatory synapses equally.

Because cortex- and thalamus-derived presynaptic terminals can be visualized by VGLUT1- and VGLUT2-positive signals, respectively^17^, we counted the numbers of VGLUT1- or VGLUT2-positive excitatory synapses from the control or DAPS dorsal striatum to determine which subset of excitatory inputs was excessive due to the inhibition of astrocytic phagocytosis. Although there was no change in the number of VGLUT2- or VGLUT2+PSD-95-positive synapses in the DAPS group (Fig. 1e, f, i, l, Supplementary Fig. 2a), our results revealed a significant increase in VGLUT1 density (Fig. 1e-h). Additionally, there was a trend toward an increase in VGLUT1+PSD-95- positive synapses in DAPS mice compared to control mice (Fig. 1e, f, k, Supplementary Fig. 2a). These results suggest that a subpopulation of excitatory synapses in the dorsal striatum, such as corticostriatal synapses, is preferentially targeted and phagocytosed by astrocytes.

To directly test the preferential phagocytosis of cortex-derived axons over thalamus- originated axons, the amount of engulfed presynaptic materials in the astrocytic cytoplasm was evaluated. To label cortex- or thalamus-originating axons in the dorsal striatum, an Alexa 568-labeled dextran tracer was injected into representative cortical or thalamic areas innervating the dorsal striatum, such as the primary/secondary motor cortex (M1/M2) or thalamic parafascicular nuclei/central lateral (PF/CL), respectively (Fig. 2a; Supplementary Fig. 3). Our results showed that the number of cortical axons engulfed was greater than the number of thalamic axons in the control striatum (Fig. 2d-e). However, the DAPS mouse striatum exhibited reduced engulfment of cortical tracers, but the engulfment of thalamic tracers was unchanged (Fig. b-e). Selective engulfment of astrocytic phagocytosis was also demonstrated through 3-dimensional (3D) reconstruction images (Supplementary Fig. 4). Some cortical terminals were still engulfed despite the impairment of MEGF10/MERTK-dependent phagocytosis, possibly because astrocytes express various mechanisms for engulfment beyond MEGF10/MERTK-dependent signaling, such as AXL receptor tyrosine kinase and brain-specific angiogenesis inhibitor-1 (BAL1)^18,19^.

**Figure 2.**
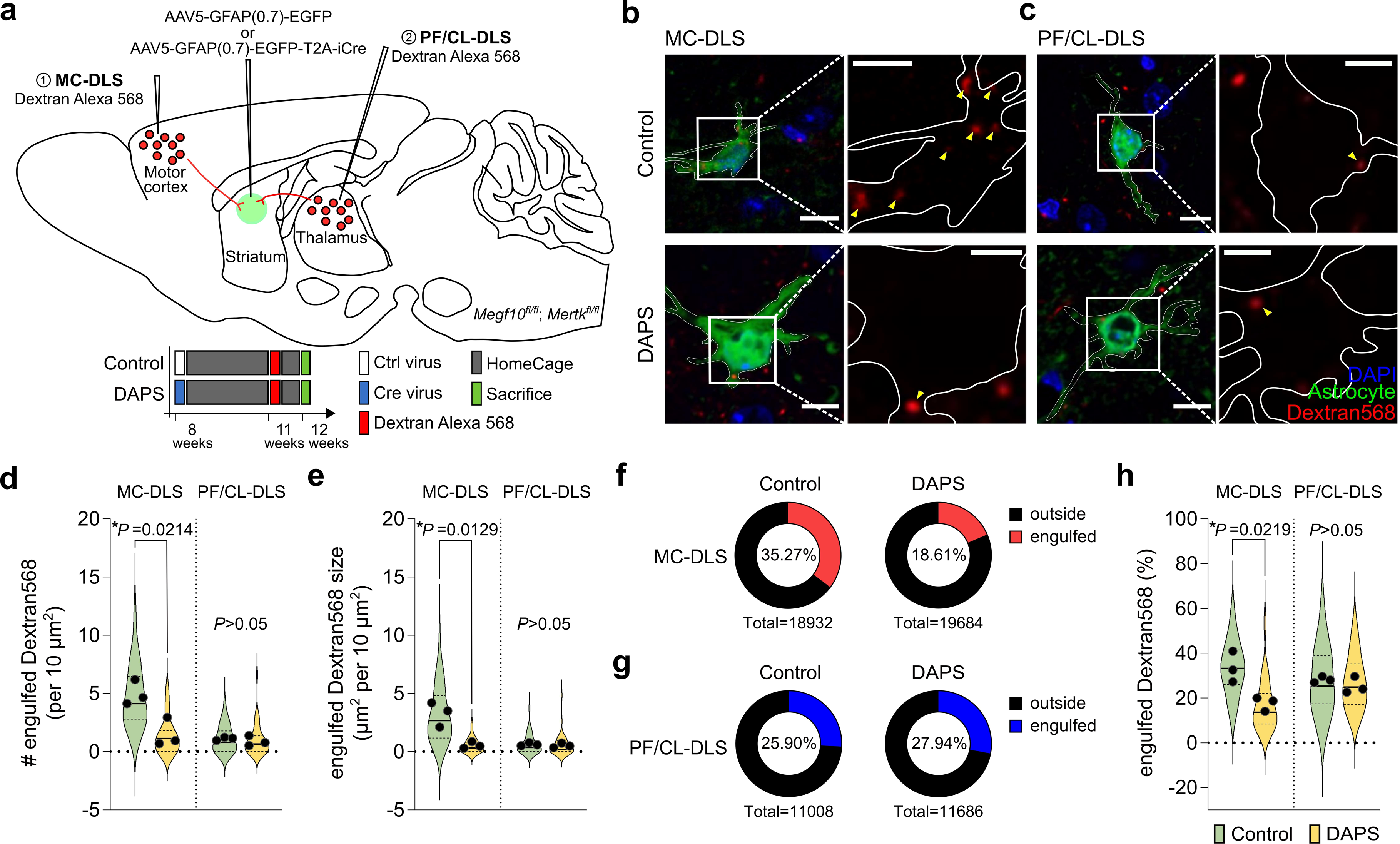
Corticostriatal synapses are preferentially phagocytosed by striatal astrocytes. **a.** Schematic diagram showing the phagocytosis assay using a fluorescent tracer (Alexa 568-conjugated dextran). **b-c.** Representative confocal images (left) and expanded panels (right) of astrocytes (green; EGFP) with fluorescent tracers (red; Dextran568) derived from the motor cortex (MC-DLS, **b**) or thalamus (PF/CL-DLS, **c**) in the striatum of control and DAPS mice. The white outlines show territories of the observed astrocytes. Scale bars= 10 μm. Scale bars= 5 μm in the expanded panels. **d-e.** Summary of the number (**d**) and size (**e**) of phagocytosed fluorescent tracers found in the astrocytic cytosol in control and DAPS mice. MC-DLS: control, n= 39 images from 3 mice; DAPS, n= 32 images from 3 mice. PF/CL-DLS: control, n= 31 images from 3 mice; DAPS, n= 34 images from 3 mice. **f.** Percentages of engulfed fluorescent tracers among all cortex-derived axonal tracers. **g.** Percentages of engulfed florescent tracers among all thalamus-derived axonal tracers. **h.** Summary of the percentage of engulfed fluorescent tracers out of all cortex- or thalamus-derived axonal tracers. MC-DLS: control, n= 34 images from 3 mice; DAPS, n= 40 images from 3 mice. PF/CL-DLS: control, n= 38 images from 3 mice; DAPS, n= 41 images from 3 mice. Unpaired student’s t-test. Mean ± s.e.m.

Next, when comparing the percentage of engulfed tracers among total M1/M2 or PF/CL-derived tracers, the DAPS mouse striatum only showed a significant reduction in phagocytosing M1/M2-derived tracers (Fig. 2f-h). Together, these data indicate that striatal astrocytes preferentially phagocytose cortical inputs over thalamic inputs.

### Preferential regulation of corticostriatal activity by astrocytic synapse phagocytosis

Because our results showed that cortex-derived synapses were preferentiall phagocytosed by striatal astrocytes (Figs. 1 and 2), we next asked whether inhibition of astrocytic phagocytosis affects corticostriatal synaptic transmission. To evoke input- specific synaptic transmission in the striatum, selective excitation of cortical and thalamic presynaptic fibers was achieved by introducing AAV containing hSyn promoter-driven Chronos (AAV-hSyn-Chronos-tdTomato) into M1/M2 and PF/CL, respectively^20,21^. The blue light (470 nm)-elicited excitatory postsynaptic potentials (EPSPs) was then recorded from MSNs of control or DAPS slices (Fig. 3a, b, Supplementary Fig. 5a).

**Figure 3.**
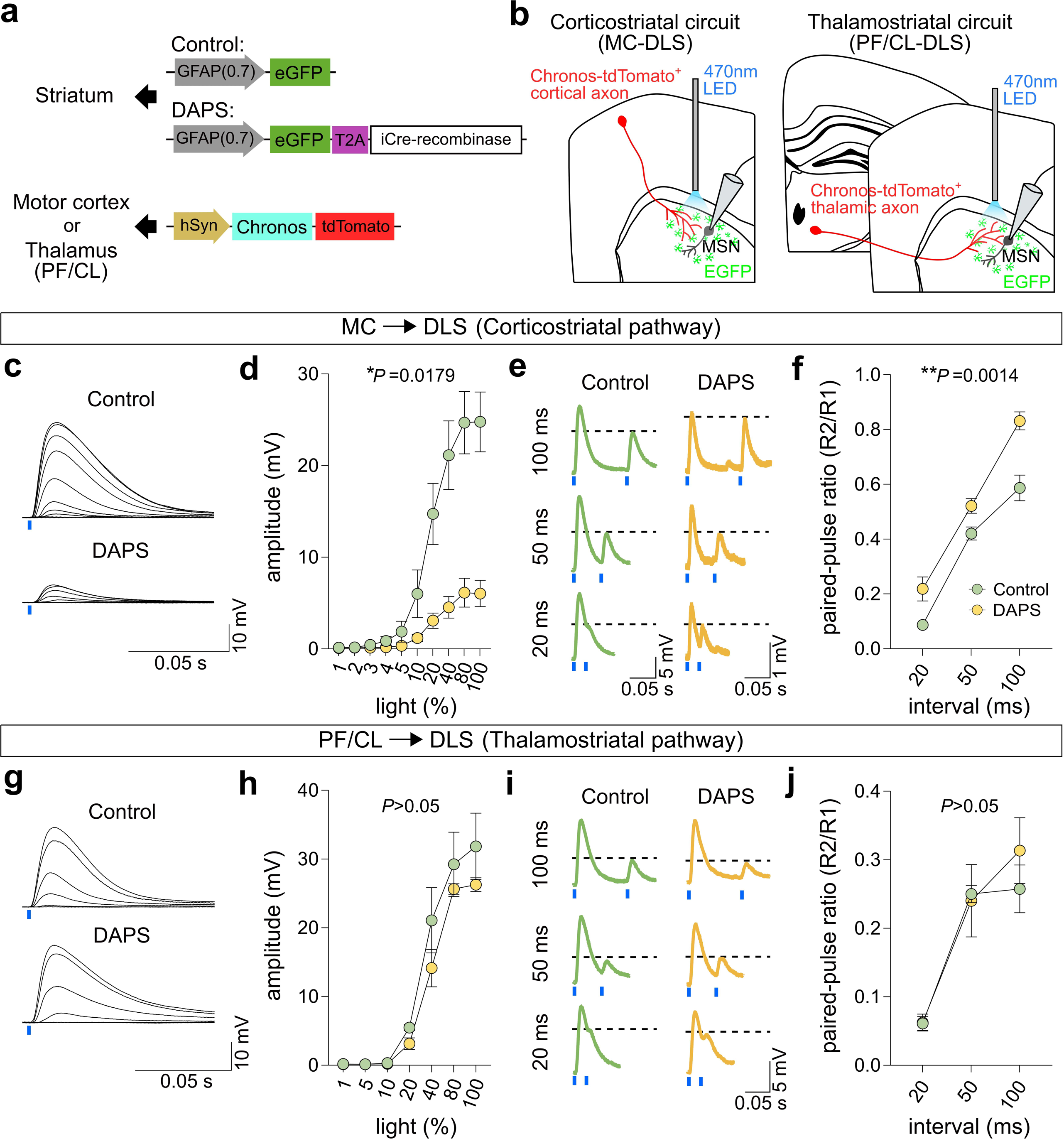
Astrocytic phagocytosis selectively regulates corticostriatal transmission. **a.** Illustration of AAVs injected into the motor cortex or thalamus in control and DAPS mice. **b.** Schematic diagram showing optogenetic MC-DLS (left) or PF/CL-DLS synaptic transmission (right). **c.** Representative traces showing optogenetic MC-DLS EPSPs evoked by illumination with blue light (470 nm) at intensities ranging from 1∼100% in control or DAPS slices. **d.** Dot plots depicting the average responses of light-evoked MC-DLS EPSPs recorded from control or DAPS striatal slices. Control, n= 18 cells from 3 mice; DAPS, n= 17 cells from 3 mice. **e.** Representative traces showing the paired-pulse ratio (PPR) of optogenetic MC-DLS in control or DAPS slices. **f.** Dot plots depicting the average MC-DLS PPR in control or DAPS-treated striatal slices. Control, n= 17 cells from 3 mice; DAPS, n= 13 cells from 3 mice. **g.** Representative traces showing optogenetic PF/CL-DLS EPSPs evoked by illumination with blue light (470 nm) at intensities ranging from 1∼100% in control or DAPS slices. **h.** Dot plots depicting the average responses of light-evoked PF/CL-DLS EPSPs recorded from control or DAPS striatal slices. Control, n= 12 cells from 3 mice; DAPS, n= 11 cells from 3 mice. **i.** Representative traces showing the paired-pulse ratio (PPR) of optogenetic PF/CL-DLS in control or DAPS slices. **j.** Dot plots depicting the average PF/CL-DLS PPR from control or DAPS striatal slices. Control, n= 12 cells from 3 mice; DAPS, n= 11 cells from 3 mice. The blue bars indicate the timing of blue light illumination. Repeated measures two-way ANOVA test. Mean ± s.e.m.

First, we tested whether light-evoked EPSPs (optogenetic EPSPs) represent monosynaptic and glutamatergic synaptic responses via cortico- or thalamostriatal pathways (Supplementary Fig. 5). Our results showed that cotreatment with bicuculline, 4-aminopyridine (4-AP), and tetrodotoxin (TTX) did not affect light-evoked cortico- or thalamostriatal EPSPs (Supplementary Fig. 5b, c), indicating that optogenetic EPSPs are mediated by monosynaptic transmission. These optogenetic EPSPs are also verified to be glutamatergic transmission, because additional treatment with D-2-amino-5- phosphonopetanoate (D-APV) and 2,3-dioxo-6-nitro-7-sulfamoyl-benzo[f]quinoxaline (NBQX) abolished these light-evoked EPSPs (Supplementary Fig. 5b, c).

Under these conditions, we next elevated cortical or thalamic axon firing by increasing the light intensity and measured the resulting EPSP responses (Fig. 3). These manipulations resulted in an exponential growth and saturation of optogenetic EPSP amplitudes in both the corticostriatal and thalamostriatal pathways (Fig. 3c, d, g, h). However, the increase in corticostriatal optogenetic EPSPs was significantly delayed in DAPS slices (Fig. 3c, d), while thalamostriatal optogenetic EPSPs were not influenced (Fig. 3g, h). In addition, the paired-pulse ratio (PPR) at corticostriatal synapses was significantly increased in DAPS slices (Fig. 3e, f), with no alterations in the thalamostriatal PPR (Fig. 3i, j), suggesting the involvement of reduced presynaptic release in abnormal corticostriatal activity in the DAPS mice. Together, these results support our findings that astrocytic phagocytosis preferentially targets corticostriatal synapses in the dorsal striatum.

### Requirement of astrocytic phagocytosis for corticostriatal synaptic plasticity

Because our data showed that astrocytic phagocytosis selectively modulates corticostriatal synapses in the striatum, we next tested whether N-methyl-D- aspartate (NMDA) receptor-dependent corticostriatal plasticity, which is essential for motor and procedural learning^15,22–25^, also depends on astrocytic phagocytosis.

Corticostriatal long-term potentiation (LTP) was elicited by illuminating theta-burst stimulation (TBS) with patterned blue light (optogenetic TBS; see Methods)^20^ in Chronos-expressing cortical afferents in control or DAPS slices. Consistent with a previous report^20^, optogenetic TBS in control slices was sufficient for inducing NMDAR-dependent potentiation of corticostriatal EPSPs lasting longer than 60 minutes (Fig. 4a, c). However, in the DAPS slices, optogenetic TBS-induced corticostriatal LTP was significantly diminished (Fig. 4a, c). These results indicate that long-term plasticity of the corticostriatal pathway depends on synapse reorganization by astrocytic phagocytosis.

**Figure 4.**
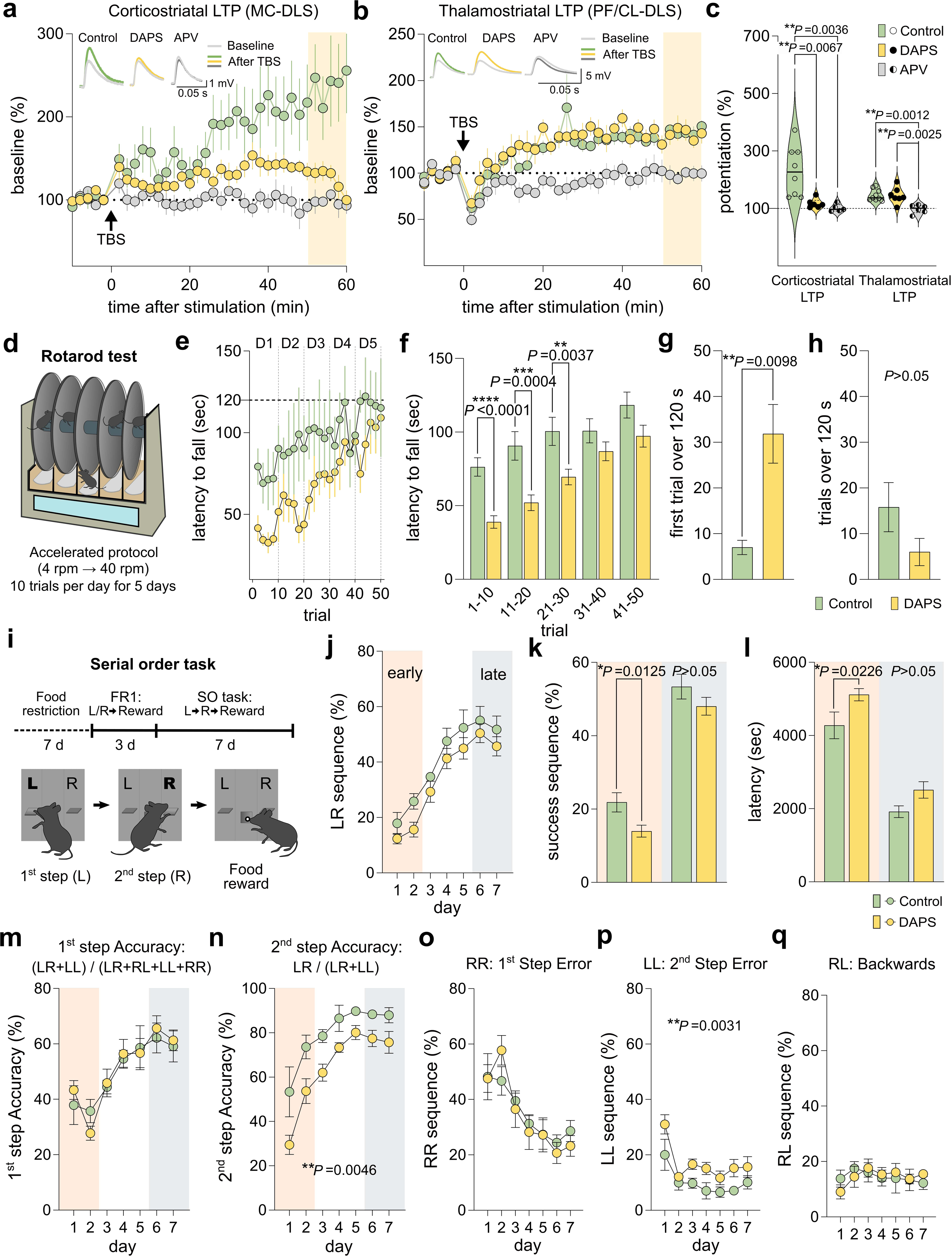
The astrocytic MEGF10/MERTK complex is required for normal corticostriatal plasticity and learning. **a.** Summary of relative changes in the amplitude of optogenetic MC-DLS EPSPs before and after optogenetic TBS at the indicated time points (black arrow). Insets: representative optogenetic MC-DLS EPSPs at baseline (gray) or 50∼60 min after TBS (control: green; DAPS: yellow; APV: dark gray). **b.** Summary of relative changes in the amplitude of the optogenetic PF/CL-DLS EPSPs before and after the administration of optogenetic TBS at the indicated time points (black arrow). Insets: Insets: representative optogenetic PF/CL-DLS EPSPs at baseline (gray) or 50∼60 min after TBS (control: green; DAPS: yellow; APV: dark gray). **c.** Left: Bar graphs depicting average potentiated optogenetic MC-DLS EPSPs 50∼60 min after TBS (yellow boxes in a). Control, n= 8 cells from 5 mice; DAPS, n= 6 cells from 5 mice; APV, n= 5 cells from 3 mice. Right: average potentiated optogenetic PF/CL-DLS EPSPs 50∼60 min after TBS (yellow boxes in b). Control, n= 9 cells from 4 mice; DAPS, n= 7 cells from 4 mice; APV, n= 6 cells from 3 mice. One-way ANOVA with Tukey’s multiple comparisons test. **d.** Schematic diagram of the rotarod test. **e.** Latency to fall during rotarod task. Note that 10 trials per day were presented as 5 dots by averaging every two trials. **f.** Bar graphs depicting the average latency to fall during rotarod learning. **g.** Bar graphs depicting average trial it takes for mice to stay on the rotarod for over 120 seconds during their initial attempt. **h**. Bar graphs depicting average number of trials showing mice stayed on the rod over 120 seconds. Control: n= 5 mice; DAPS: n= 7 mice. **i.** Schematic diagram of the SO task. **j.** Percentage of correct left-right (LR) sequences among total attempts (LL+LR+RL+RR) during SO training. The early phase (orange) and late phase (gray) of learning are indicated. **k.** Bar graphs depicting the average percentages of success LR sequences during the early and late periods of learning. **l.** Bar graphs indicating the average time to reach completion for the early and late periods of learning. **m.** Average percentage of first-step accuracy (LR + LL) among all trials during SO training. **n.** Average percentage of second step accuracy (LR) in trials beginning with the correct first sequence (LL + LR) during SO training. **o-q.** Percentage of sequences completed in each potential order. RR: first step error sequences (**o**); LL: second step error sequences (**p**); RL: backward sequences (**q**). Control: n= 5 mice; DAPS: n= 10 mice. Control: n= 5 mice; DAPS: n= 10 mice. Repeated measure Two-way ANOVA test. Bar graphs were analyzed using unpaired student’s t-test. Error bars: SEM.

On the other hand, astrocytic phagocytosis was not involved in thalamostriatal LTP because NMDAR-dependent optogenetic TBS-LTP was still observed either from control or DAPS striatal slices expressing Chronos at thalamic afferents (Fig. 4b, c). These results suggest that thalamostriatal plasticity may require mechanisms other than astrocytic MEGF10/MERTK-mediated phagocytosis for circuit connectivity and plasticity.

### Contribution of astrocytic phagocytosis-mediated synapse turnover to motor sequence learning

We next tested whether the preferential regulation of the corticostriatal pathway by astrocytic phagocytosis contributes to striatal learning and memory. Control and DAPS mice were subjected to the rotarod test (Fig. 4d) and the serial order (SO) task (Fig. 4i), both of which are known to depend on the corticostriatal pathway^23,26^.

Both control and DAPS mice showed a progressive learning curve during the accelerated rotarod task (Fig. 4d), with 10 trials conducted each day over a total of 5 days. Although there was no significant difference in the overall trend of the learning curve (Fig. 4e), we found that DAPS mice demonstrated a significantly lower performance level of motor skill learning than control mice during the first three days (Fig. 4f). However, DAPS mice eventually reached normal levels in the late phase of learning (Fig. 4f). When comparing the trial at which each group first reached the average learning point of the control group on day 5 (120 seconds), DAPS mice took more trials to reach this desired learning point than control mice (Fig. 4g). However, when assessing the overall trials that exceeded the desired learning point, no difference between the two groups was found (Fig. 4h). These results suggest that astrocytic refinements of corticostriatal pathway through phagocytosis mechanisms are important for early periods of motor skill learning.

Next, to test whether another corticostriatal pathway-dependent procedural learning also requires the role of astrocytic phagocytosis, the SO task was conducted. Prior to SO training, the mice were subjected to food restriction for 7 days and then trained on a fixed ratio one (FR1) schedule for 3 days to reinforce the correlation between the lever and food reward (Fig. 4i). During this period, comparable weight loss, normal locomotion and anxiety, and similar amounts of food consumption were observed in both control and DAPS mice (Supplementary Fig. 6, 7 and 8a-c), indicating that astrocytic synapse phagocytosis does not affect motivation for food reward or the striatal circuit involved in general motor behaviors.

After the FR1 schedule, we introduced the SO task for testing motor sequence learning for 7 days. A food reward was delivered only after the completion of the lever pressing sequence (left → right; LR) (Fig. 4i). The control mice showed a progressive increase in the percentage of successful LR sequences over the course of training (Fig. 4j). Similar to the rotarod results (Fig. 4e), there was no significant difference in the overall learning curves between control and DAPS mice (Fig. 4j). However, DAPS mice exhibited a significantly lower percentage of the successful LR sequence during the early period (Day 1∼2; Fig. 4j, k). Moreover, the average time to completion of the SO task was also delayed during the early learning period in the DAPS group (Fig. 4l). After the early learning period, DAPS mice exhibited a comparable success rate to control mice (Day 6∼7; Fig. 4j, k), indicating that DAPS mice could achieve the desired learning goal in the late learning period as control mice. Because both the overall response rate and food consumption remained consistent throughout the SO task sessions (Supplementary Fig. 8), delayed learning of the SO task during the early phase was not caused by alterations in motivation due to the inhibition of astrocytic phagocytosis.

Reduced learning in the early phase of the SO task was prominent when the sequential accuracy of the SO task was compared. Among the total correct sequences (LRs) that began with a correct initial step (L) and subsequently completed with a correct second step (R), the DAPS mice displayed reduced accuracy of second step (Fig. 4n), with comparable accuracy in the first step with control group (Fig. 4m). Indeed, DAPS mice exhibited a significantly increased level of adopting the LL sequences, representing elevated second-step error, instead of the correct LR sequence (Fig. 4o-q). Together, these results indicate that astrocytic maintenance of the corticostriatal circuit contributes to the early phase of motor skill and sequence learning in adult mice.

## Discussion

In this study, we demonstrated that astrocytic MEGF10/MERTK consistently regulate synapse reorganization in the adult striatum. These results support the notion that striatal and hippocampal astrocytes eliminate unnecessary synapses via the same phagocytosis mechanisms dependent on MEGF10/MERTK. Given the characteristics of the hippocampus and striatum, astrocytes likely contribute to regulating neural circuit connectivity in multiple brain areas, where rapid turnover and repatterning of neural circuits are necessary for experience-dependent synaptic plasticity.

Similar to astrocytes in the adult hippocampal CA1 area^9^, striatal astrocytes selectively target and phagocytose excitatory synapses (Fig. 1). Given the recent report that microglial MERTK regulates inhibitory postsynapses^27^, astrocytes and other glial cells may differentially recognize and phagocytose synapses for circuit homeostasis. Indeed, the mechanism by which astrocytes recognize specific synapse types are unclear, but two possibilities exist: recognition of structural differences in synapses or detection of ’eat-me’ signals on target cells^28–31^. Astrocytic phagocytosis primarily targets excitatory synapses, particularly MC-derived corticostriatal synapses rather than PF/CL-derived thalamostriatal synapses. Excitatory synapses and MC-derived synapses mainly form on dendritic spines, while inhibitory synapses and PF/CL-derived thalamostriatal synapses typically target dendritic shafts. Such resemblance in astrocytic targeting of synapse phagocytosis raises the possibility that specific synaptic structures determine the selectivity of astrocytic phagocytosis. However, further analysis of ventral anterior/lateral nucleus (VA/VL)-derived thalamostriatal synapses forming dendritic spine structures in the striatum^32^ revealed no alteration in astrocytic engulfment of VA/VL-derived terminals by *Megf10/Mertk* knockout (Supplementary Fig. 9). It is thus unlikely that astrocytes recognize structural features of synapses to determine the target of phagocytosis.

Instead, the involvement of ‘eat-me’ signals emerge as a more plausible mechanism for astrocytic phagocytosis. To date, the molecular identity of synaptic "eat-me" signals directing synapse- or circuit specific phagocytosis via MEGF10 and MERTK remains to be fully understood. In the immune system, MERTK has been shown to bind to phosphatidylserine (PS), a well-known eat-me signal that is exposed on the surface of apoptotic cells^33^. MEGF10 can also recognize PS indirectly through linking molecules like C1q and TTR-52^34,35^. Alternately, MEGF10 can recognize other eat-me signals than PS, such as endoplasmic reticulum proteins like Pretaporter or lipoteichoic acid for phagocytosis^36,37^. Therefore, a variety of potential ligands for MEGF10 and MERTK may exist in the brain, and future studies will be necessary to investigate the molecular identity and mechanisms associated with the recognition of this specific type of synapses.

In particular, our results showed that astrocytic phagocytosis is biased toward corticostriatal synapses rather than thalamostriatal synapses (Figs. 1, 2). This result provides the first evidence that astrocytes recognize a specific type of excitatory synapse among synapses originating from diverse input areas. Although the underlying mechanisms by which striatal astrocytes can recognize certain types of synapses are unclear, it is possible that the molecular heterogeneity of diverse types of excitatory synapses^38–40^ is responsible for the input specificity of astrocytic synapse phagocytosis.

Our results showed that the abnormal corticostriatal transmission in the DAPS striatum is caused by the homeostatic downregulation of presynaptic functions. DAPS mice exhibited abnormal phenotypes in striatal synapses, characterized by decreased optogenetically evoked EPSC amplitude and PPR (Fig. 3), as well as increased VGLUT1(+) excitatory synapse number (Fig. 1) and sEPSC frequency (Fig. 1). These alterations resemble those observed in hippocampal CA3-CA1 synapses following the knockout of the astrocytic *Megf10* gene^9^. In both cases, reduced astrocytic synaptic phagocytosis led to excessive excitatory connectivity, prompting a compensatory decrease in presynaptic release to mitigate excessive excitatory network activity. We suggest that the abnormal synaptic properties of DAPS mice result from the homeostatic suppression of cortical presynaptic release. Consequently, the inhibition of astrocytic phagocytosis led to impairments in long-term synaptic plasticity in both the hippocampus^9^ and striatum (Fig. 4). Together, these results suggest that constant excitatory synapse turnover by astrocytes is crucial for synapse connectivity optimized for experience-dependent plasticity.

We showed that astrocytic synapse reorganization significantly contributed to the early phase of motor sequence learning (Fig. 4). This is probably because the motor cortex becomes independent in the later stage of motor learning^41^, and motor skill learning and execution are distinctly regulated by corticostriatal and thalamostriatal pathways, respectively^15^. Therefore, the early learning delay in DAPS mice may result from excessive corticostriatal circuit activity and deficits in plasticity. The motor performance in DAPS mice appears to be normal in the late phase of learning, probably due to the reduced dependency of motor performance on corticostriatal activity during the late phase or compensatory involvement of other striatal circuits not associated with astrocytic phagocytosis.

In conclusion, our finding of preferential regulation of the corticostriatal pathway by astrocytic phagocytosis supports the idea that astrocytes preferentially regulate a specific subset of excitatory synapses in the mature brain. Further studies will be needed to identify other mature brain circuits dependent on astrocytic synapse reorganization and molecular signaling directing the circuit specificity of astrocytic phagocytosis.

## Acknowledgments

We thank all members of the Park laboratory for helpful discussion. This work was supported by KBRI basic research program through Korea Brain Research Institute funded by Ministry of Science and ICT [24-BR-01-02 (H.P.)]. Instruments and whole- cell patch clamp data were acquired at Brain Research Core Facilities in KBRI.

## Author contributions

J.K., W.S.C. and H.P. designed the projects. J.K. performed all virus and tracer injection experiments. J.K. performed and analyzed all electrophysiology, optogenetic recording, behavior, and phagocytosis assay experiments. H.K. performed and analyzed synapse quantification experiments. H.P. supervised the project. J.K. and H.P. wrote the paper.

## Methods

### Animals

All mouse experiments conformed to Institutional Animal Care and Use Committee (IACUC) protocols of the Korea Brain Research Institute (KBRI). We followed all proper ethical regulations. LoxP-floxed *Megf10* (*Megf10^fl/fl^*) and loxP-floxed *Mertk* (*Mertk^fl/fl^*) mice were generated by the Stanford Transgenic, Knockout and Tumor Model Center (TKTC). *Megf10^tm^*^1^*^(KOMP)Vlcg^* (straight *Megf10* knockout, st*Megf10* KO) and *Mertk^tm^*^1^*^(KOMP)Vlcg^* (straight *Mertk* knockout, st*Mertk* KO) mice were obtained from Ben Barres’ laboratory^7,9^. The *Megf10^fl/fl^* and *Mertk^fl/fl^* lines were crossed together to produce double loxP-floxed *Megf10* and *Mertk* (*Megf10^fl/fl^*;*Mertk^fl/fl^*) mice. The mouse lines were maintained through crossing with C57BL/6 mice in a standard plastic cage. A 12-hour light/12-hour dark cycle was implemented, and the temperature inside the cage was maintained between 20 °C and 23 °C. All experiments involving mutant mice were performed blindly with other littermates. All mice were randomly assigned to groups for experimentation.

### Stereotaxic injection

Mice were anesthetized with isoflurane (Hana Pharm) in a sealed acrylic box and maintained under a low-flow anesthesia delivery system (SomnoSuite). For deletion of astrocytic phagocytosis receptors, either AAV5::GFAP (0.7)-Cre-eGFP-T2A-iCre- WRPE or AAV5::GFAP (0.7)-eGFP-WPRE (6.4x10^12^GC/ml and 1.2x10^13^GC/ml, respectively, purchased from Vector Biolabs) was stereotaxically injected bilaterally into the dorsal striatum (ML: ± 1.90 mm, AP: +0.79 mm from bregma, DV: -2.70 mm from the brain surface) of 7-9-week-old *Megf10^fl/fl^;Mertk^fl/fl^* mice. For the phagocytosis assay, an Alexa 568-labeled dextran tracer was stereotaxically injected bilaterally into either the motor cortex (ML: ± 0.75 mm, AP: +1.97 mm from bregma, DV: -0.50 mm from the brain surface), thalamic central lateral (CL; ML: ± 0.81 mm, AP: -1.50 mm from bregma, DV: -2.94 mm from the brain surface)/parafascicular nuclei (PF; ML: ± 0.61 mm, AP: -2.30 mm from bregma, DV: -3.34 mm from the brain surface) or ventral anterior/lateral nucleus (VA/VL; ML: ± 1.32 mm, AP: -0.83 mm from bregma, DV: - 3.30 mm from the brain surface) of control or DAPS mice, respectively. For optogenetic recording, AAV2/2::hSyn-Chornos-tdTomato (2x10^12 GC/ml, purchased from BrainVTA) was injected into the motor cortex or thalamic PF/CL simultaneously with AAV injection for control and DAPS production. All surgeries were performed using a stereotaxic frame (Kopf) and a Nanojector III (Drummond). A glass pipette needle (WPI) was used for the nanoinjector, and pulling of the glass pipette was performed using a motorized pipette puller (Sutter instrument). After injection, the incision on the head was closed with wound closure clips (Alzet). After the surgery, the mice were allowed to recover in a heated cage for an hour before being returned to their home cage.

### Immunohistochemistry

Mice were deeply anesthetized with 1-2% isoflurane and intracardially perfused with 1X PBS (Welgene), followed by ice-cold 4% paraformaldehyde (PFA, BIOSESANG) in PBS. Mice brains were isolated and post-fixed overnight in PFA at 4 °C, and then dehydrated in 30% sucrose in PBS for 48 hours. The dehydrated brains were embedded in OCT compound (Leica) and frozen at -80 °C. Forty-micrometer brain slices were collected using cryostat microtomes (Leica). The sections were blocked with 10% goat serum and 0.2% Triton X-100 in 1× PBS for 1 h at room temperature (RT) and incubated overnight at 4 °C with primary antibodies diluted in the blocking solution. The primary antibodies used were as follows: guinea pig anti-VGLUT1 (#AB5905, Millipore, 1:2000 dilution), chicken anti-VGLUT2 (#135 416, Synaptic Systems, 1:500 dilution), rabbit anti-PSD95 (#3450, Cell Signaling, 1:200 dilution), guinea pig anti- VGAT (#131 004, Synaptic Systems, 1:500 dilution), rabbit anti-Gephyrin (#147 008, Synaptic Systems, 1:500 dilution), guinea pig anti-S100B (#287 004, Synaptic Systems, 1:500 dilution), rabbit anti-Iba1 (#019-19741, FujiFilm, 1:500 dilution), rabbit anti- MEGF10 (#ABC10, Millipore Sigma, 1:200 dilution) or rabbit anti-MERTK (#Ab216564, Abcam, 1:200 dilution). The sections were then washed with 0.2% Triton X-100 in 1×PBS (PBST) and treated with appropriate secondary antibodies conjugated with Alexa Fluor in blocking solution for 2 h at RT. The following secondary antibodies were used: goat anti-rabbit Alexa Fluor 405 (#A31556, Invitrogen, 1:200 dilution), goat anti-guinea pig Alexa Fluor 546 (#A11074, Invitrogen, 1:200 dilution), goat anti- chicken Alexa Fluor 647 (#A21449, Invitrogen, 1:200 dilution), goat anti-rabbit Alexa Fluor 647 (#A21245, Invitrogen, 1:200 dilution), donkey anti-guinea pig Alexa Fluor 647 (#706-605-148, Jackson ImmunoResearch, 1:200 dilution) or goat anti-rabbit Alexa Fluor 568 (1:200, Cat #11011, 1:200). The stained sections were washed with PBST and mounted on adhesive-coated glass slides (MARIENFELD) with Vectashield Hardset Antifade Mounting Medium (H-1400, Vector Lab). Images were acquired using confocal microscope (Nikon, A1 Rsi/Ti-E or Leica, STELLARIS 8) or slide scanner (3DHISTECH, Pannoramic scans).

### Phagocytosis assay

Confocal images of the dorsal striatum sections were acquired using a Nikon A1 Rsi/Ti-E (60x oil immersion optical lens) for quantification, as described below. Z-stack imaging was employed to examine 4 to 6 astrocytic cell bodies and their extensions in a 3D space for the dextran engulfment test. A single image from the captured volumes of 215 um x 215 um x 30 ∼ 35 um dimensions featuring an astrocyte cell body was selected to count the engulfed dextran puncta. To analyze the amount of Alexa 568-labeled dextran engulfed within the astrocyte cytoplasm, the EGFP^+^ astrocyte processes were visualized, and the number of engulfed puncta was measured. The data generated were used to calculate the density of the engulfed dextran puncta (number of engulfed dextran puncta/area of EGFP^+^ astrocytes), the size of the engulfed dextran puncta (size of engulfed dextran puncta/area of EGFP+ astrocytes), and the percentage of engulfed dextran puncta (number of engulfed dextran puncta/total projected dextran puncta) in the corticostriatal or thalamostriatal pathways. The colocalization assay was conducted using the DiAna plugin^42^. For 3D reconstruction of post-processed stacked confocal images by LAS X Lightning, IMARIS (Bitplane) was utilized.

### Quantification of synapse numbers

Confocal images of excitatory synaptic compartments in the dorsal striatum were acquired using a Leica STELLARIS 8 (63x oil immersion optical lens), and images of inhibitory synaptic compartments were obtained using a Nikon A1 Rsi/Ti-E (60x oil immersion optical lens). To avoid biased interpretation due to inconsistent selection of image fields (e.g., cell bodies or processes), the analysis was conducted on a single confocal optical section (optical section = 0.896 µm) containing at least 4 to 6 complete astrocyte cell bodies with processes. At least 20 confocal optical sections per mouse were analyzed to ensure reliable data acquisition. All channels, including the presynaptic and postsynaptic compartments, were separated using ImageJ software. The number of excitatory/inhibitory synaptic puncta (VGLUT1&2, VGLUT1, VGLUT2, VGAT, PSD95, or gephyrin) was counted using the Analyze Particles function in ImageJ software. The number of colocalized puncta (pre- and postsynaptic together; VGLUT1+PSD95, VGLUT2+PSD95, or VGAT+gephyrin) was measured using the DiAna plugin^42^.

### Whole-cell patch clamp

For whole-cell patch clamp recordings, acute brain slices were obtained from 12- to 16- week-old mice. The standard artificial cerebral spinal fluid (ACSF) consisted of (in mM) 124 NaCl, 2.5 KCl, 1.2 NaH_2_PO_4_, 24 NaHCO_3_, 5 HEPES, 2 CaCl_2_, 2 MgCl_2_, and 13 glucose (pH 7.3). Mice were deeply anesthetized with isoflurane and intracardially perfused with ∼20 ml of slicing ACSF containing (in mM) 93 N-methyl-D-glutamine (NMDG)-Cl, 93 HCl, 2.5 KCl, 1.2 NaH_2_PO_4_, 30 NaHCO_3_, 20 HEPES, 5 sodium ascorbate, 2 thiourea, 3 sodium pyruvate, 12 N-acetyl-L-cysteine (NAC), 0.5 CaCl_2_, 10 MgCl_2_, and 25 glucose (pH 7.3). Coronal slices containing the striatum (350 μm thick) were dissected using a VF-200-OZ Compresstome (Precisionary) using the slicing ACSF and recovered at 32.5 °C in recovery ACSF (in mM; 104 NaCl, 2.5 KCl, 1.2 NaH_2_PO_4_, 24 NaHCO_3_, 5 HEPES, 5 sodium ascorbate, 2 thiourea, 3 sodium pyruvate, 12 NAC, 2 CaCl_2_, 2 MgCl_2_, and 13 glucose; pH 7.3) for 1 h.

The slices were placed in a recording chamber and continuously perfused with oxygenated standard ACSF at a rate of 2-3 ml/min at RT. Whole-cell recordings were made with a Multipclamp 700B amplifier (Molecular Devices). The data were filtered at 5 kHz and digitized at 10-50 kHz. Borosilicate glass patch electrodes with a resistance of 3-5 MΩ were filled with pipette solution containing (in mM) 140 Cs- methanesulfonate, 7 NaCl, 0.2 EGTA, 2 MgCl2, 4 Mg-ATP, 0.3 Na2-GTP, 10 Na2-phosphocreatine, and 10 HEPES (pH 7.3, 290-300 mOsm) and used for recording spontaneous excitatory postsynaptic currents (sEPSCs). To record spontaneous inhibitory postsynaptic currents (IPSCs), pipette solutions containing (in mM) 140 CsCl, 7 NaCl, 0.2 EGTA, 2 MgCl2, 4 Mg-ATP, 0.3 Na2-GTP, 10 Na2-phosphocreatine, and 10 HEPES (pH 7.3, 290-300 mOsm) were used. Bicuculline (10 μM; Tocris), 2,3- dioxo-6-nitro-7-sulfamoyl-benzo[f]quinoxaline (NBQX, 10 μM; Tocris), and D-2- amino-5-phosphonopetanoate (D-AP5, 50 μM; Tocris) were added to inhibit GABAergic or glutamatergic synaptic transmission, respectively. Striatal medium spiny neurons (MSNs) were distinguished through membrane properties (Table 1) and delayed firing patterns via current injection (data not shown).

To selectively stimulate corticostriatal and thalamostriatal pathways, optogenetic induction was utilized with brain slices injected with AAV2/2::hSyn-Chronos-tdTomato in the motor cortex or thalamus. Preparation of coronal striatal slices (350 μm thick) and whole-cell recording from MSNs were performed as described above. To induce light-evoked EPEPs, a high-power LED (at 470 nm; X-Cite) was used to deliver blue light to the slice through the microscope (Nikon). This configuration could deliver blue light at ∼2.5 mW/mm^2^ over a 0.22 mm^2^ area of the recording site using a 40X objective lens. These conditions were sufficient for eliciting stable EPSPs with a light duration of 0.5-1 ms. Bicuculline (10 μM), NBQX (10 μM), D-AP5 (50 μM), 4-aminopyridine (4- AP, 100 μM; Sigma) and tetrodotoxin (TTX, 1 μM; Alomone) were added to inhibit GABAergic synaptic transmission or confirm monosynaptic/glutamatergic synaptic transmission, respectively.

The paired-pulse ratio (PPR) of the corticostriatal and thalamostriatal pathways was measured by pairing blue light-evoked stimuli to the striatum with interstimulus intervals of 20, 50 or 100 msec. Stimulus intensity was determined by constructing an input–output relationship that plotted the amplitude of light-evoked EPSPs against stimulus intensities and then adjusted to 30–40% of the maximum amplitude of light- evoked EPSPs. After at least 10 min of stable light-evoked EPSP acquisition, the PPR was measured and calculated by dividing the amplitude of the second response by that of the first response. For optogenetic induction of corticostriatal and thalamostriatal long-term potentiation (LTP), theta-burst stimulation (TBS; 10 trains of stimuli spaced at 10 s intervals, with each train containing bursts of 4 spikes at 100 Hz and repeated 10 times at 5 Hz; Park et al., Neuron, 2014) was delivered. The data for EPSP amplitudes are presented as averages over 2 min bins. TBS-induced LTP was measured as the average EPSPs at 50-60 min. All whole-cell patch clamp recording data were analyzed by using pClamp 11.1 (Axon Instruments).

### Rotarod test

An accelerating rotarod (Panlab, Harvard apparatus) was utilized as a paradigm for motor skill learning. Mice were trained with 10 trials per day for 5 days. For each trial, mice were placed on a moving rod rotating at a constant speed of 4 rpm. The speed of the rod was then gradually increased from 4 to 40 rpm over a duration of 300 seconds. This training protocol has been previously described as a reliable test for motor skill learning^43^. All rotarod test data from both animal groups passed the normality test

### Serial order task

The serial order task (SO) was performed in the operant box (Med Associates, Inc.) for each mouse. The detailed training process for the SO task was utilized with minor modifications from published protocols^23^. The operant box consisted of left (L) and right (R) levers, and a food magazine was located at the middle of the levers. For effective motor sequence learning, the mice were subjected to food restriction for 7 days prior to the first training session. First, in the fixed ratio 1 (FR1) training, the association between lever and reward was established by delivering one 14 mg of sugar pellet (Bio-Serv) after each lever response. During the FR1 session, the mice received up to 50 pellets in a 60 min session. For the SO task, the mice had to perform two distinct and sequential responses (“L” then “R”). The delivery of one sugar pellet followed the correct LR sequence, and both correct and incorrect trials were followed by an 8-s intertrial interval. Daily SO training sessions lasted for up to 90 min or until the mouse received 50 pellets. The accuracy of the first step was determined by the percentage of trials that started with a correct first step (LL or LR), while the accuracy of the second step was defined as the proportion of trials that began with a correct initial step (LL or LR) and subsequently completed with a correct second step (LR). All serial order task data from both animal groups passed the normality test

### Open field test

The open field test was performed in a square arena (nonglossy acrylic box, 300 × 300 × 280 mm, W × D × H). To start the test, the mice were placed in the center of the box and allowed to explore for 5 min. Movement was detected automatically using Noldus EthoVision 3.0 tracking software. Measurements during the test included the total distance traveled, speed, and time spent in the center/peripheral zones.

### Elevated plus maze

Mice were allowed to explore an elevated platform (50 cm above the floor) consisting of two open (30 × 6 cm) and two closed arms (30 × 6 cm with a 20 cm tall opaque wall) with a central area (6 × 6 cm). To start the test, the mice were placed in the center of the maze facing the open arm and allowed to explore for 5 min. Movement was detected automatically using SMART VIDEO TRACKING Software (Panlab). Measurements during the test included the time spent in the open arms, closed arms, and center. The maze was cleaned with 70% ethanol before each trial.

### Statistical analysis

All statistical analyses were performed using GraphRad Prism 10 with 95% confidence. Most of the imaging data and early/late behavioral data from the serial order task and rotarod test were analyzed using Student’s unpaired t-test, as these involved comparisons between two independent groups. Repeated measures Two-way ANOVA was used for light-evoked EPSPs, paired-pulse ratio, and behavioral learning curves, where time or optogenetic stimulation was a factor. In the case of LTP data comparing potentiation percentages across groups, one-way ANOVA followed by Tukey’s multiple comparisons test was applied.

Accordingly, the unpaired t-test was used to test the significant differences of sEPSC/sIPSC responses (Fig. 1c-d), astrocyte-specific AAV expression and resulted reductions in both MEGF10 and MERTK expression (Supplementary Fig. 1e-h), numbers of excitatory and inhibitory synapses (Fig. 1g-l, Supplementary Fig. 2c-e), the amount of astrocytic engulfment of MC-DLS and PF/CL-DLS pathways (Fig. 2d, e, h), and early/late learning rate in behavior tasks (Fig. 4f-h, k, l). For analyzing corticostriatal or thalamostriatal LTP, the one-way ANOVA test was utilized to compare the percentage of potentiation across groups (Fig. 4c). The repeated two-way ANOVA test was employed to test for statistical differences in the optogenetic corticostriatal or thalamostriatal EPSPs evoked by ranged light intensities (Fig. 3d, h), paired-pulse ratios evoked by different inter-stimulus intervals (Fig. 3f, j), and behavioral data during multiple days (Fig. 4e, j, m-q).

To determine whether sample data (based on mouse) has been drawn from a normally distributed data population in behavior experiments, a normality test was applied to results acquired from the serial order task and rotarod test. We found that behavior data from both animal groups passed the normality test.

The data are presented as violin plots and individual dots, representing the distribution of individual cell/image and mouse data, respectively. A solid and two dotted lines in the violin plot represent the median and upper/lower quartile of the distribution of individual cell/image, respectively. Summary of results is expressed as the mean ± standard error of the mean (SEM). All statistical analyses were based on a per mouse basis.

## DATA availability

Information on the primary/secondary antibodies used in this study are in the Methods section. All data are available upon reasonable request. For further inquiries, please contact Corresponding author.

