## Supplementary Figure for "Selective regulation of corticostriatal synapses by astrocytic phagocytosis"

※ This file contains Supplementary Figure 1 ~ 9 and Table 1.

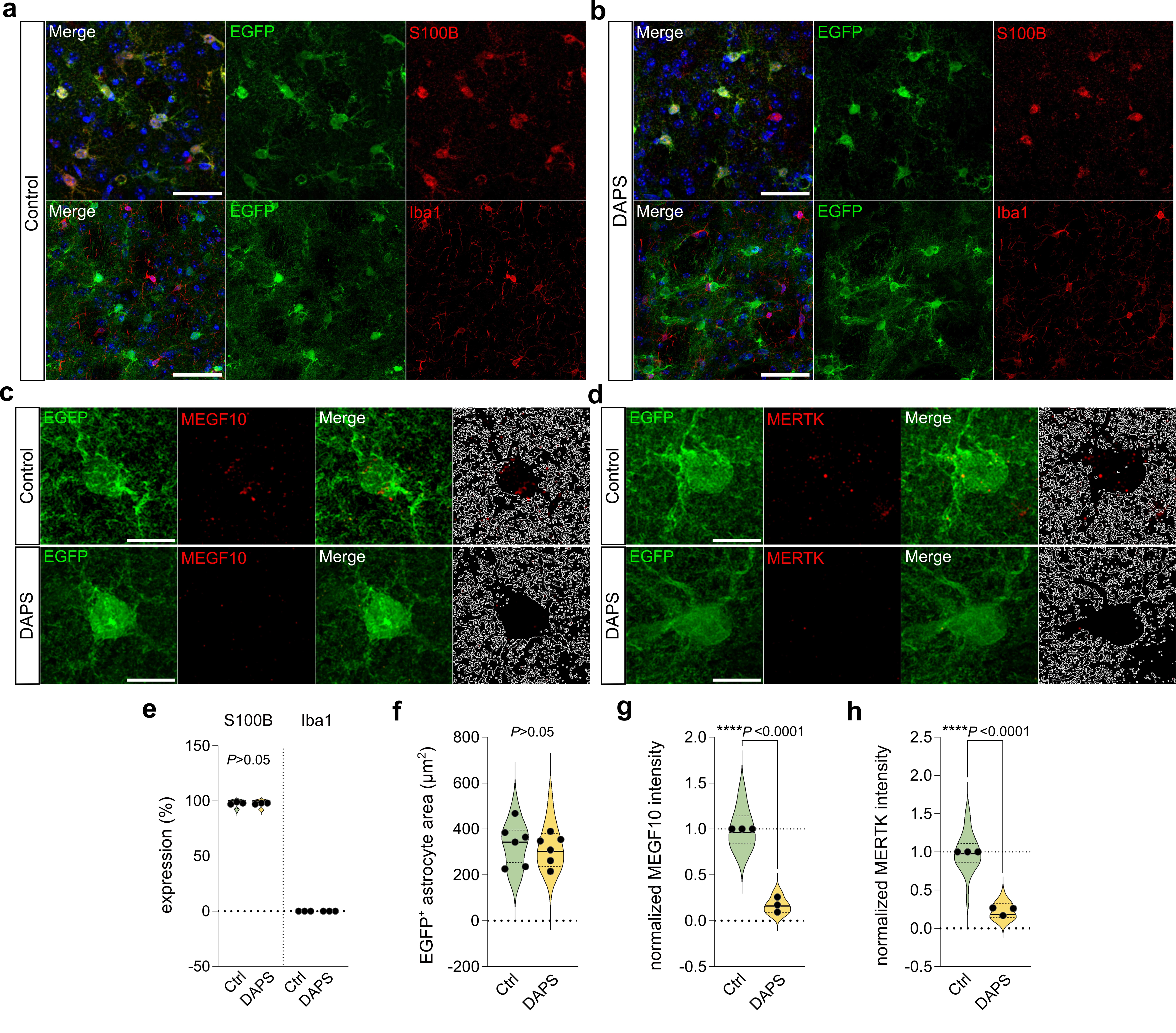

**Supplementary Fig. 1. Astrocyte-specific AAV transduction and reductions in both MEGF10 and MERTK expression in the DAPS mice**

**a-b.** Representative confocal images of astrocyte-specific expression of EGFP by AAV-GFAP0.7-EGFP (control, **a**) or AAV-GFAP0.7-EGFP-T2A-iCre (DAPS, **b**). DAPI (blue) and S100B/Iba1 (red) were indicated. scale bars= 20 μm. **c-d.** Representative confocal images of MEGF10 (**c**) or MERTK (**d**) with astrocytes in control and DAPS mice. White outlines show territories of EGFP-expressing astrocytes. Scale bars= 10 μm. **e.** Percentage of EGFP expressing cells with S100B or Iba1. S100B: Control, n= 19 images from 3 mice; DAPS, n= 20 images from 3 mice. Iba1: Control, n= 18 images from 3 mice; DAPS, n= 18 images from 3 mice. **f.** Summary of the EGFP-positive astrocyte area. **g-h.** Summary of the MEGF10 (g) or MERTK (h) intensity within EGFP-positive astrocyte area. MEGF10: Control, n= 20 images from 3 mice; DAPS, n= 20 images from 3 mice. Iba1: Control, n= 23 images from 3 mice; DAPS, n= 23 images from 3 mice. Unpaired Student’s t-test. Mean ± s.e.m.

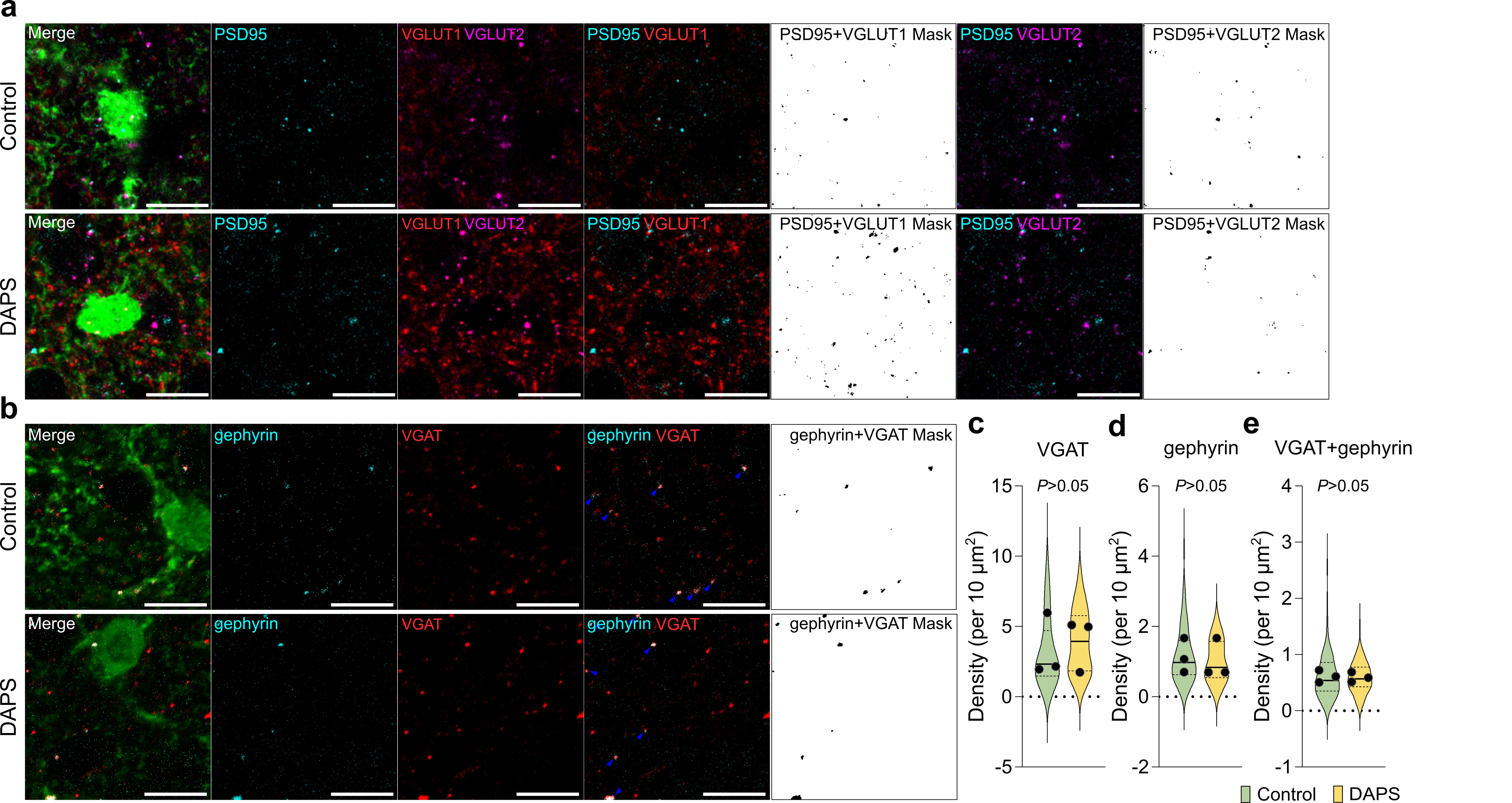

**Supplementary Fig. 2. Effects of astrocytic *Megf10/Mertk* knockout on striatal excitatory and inhibitory synapses**

**a.** Representative confocal images depicting excitatory synapse markers in the dorsal striatum of control and DAPS mice. AAV-transduced EGFP-expressing astrocytes (green), VGLUT1 (red), VGLUT2 (magenta) and PSD-95 (cyan) are shown. Binary masks showing colocalization of PSD95+VGLUT1 or PSD95+VGLUT2, respectively. Scale bars= 10 μm. **b.** Representative confocal images depicting inhibitory synapse markers in the dorsal striatum of control and DAPS mice. AAV-transduced EGFP-expressing astrocytes (green), VGAT (red) and gephyrin (cyan) are shown. Binary masks showing colocalization of gephyrin+VGAT. Scale bars= 10 μm. **c-e.** Summary of the density of VGAT (j)- and gephyrin (k)-positive particles and VGAT and gephyrin double-positive particles (l). Control, n= 77 images from 3 mice; DAPS, n= 72 images from 3 mice. Unpaired student’s t-test. Mean ± s.e.m.

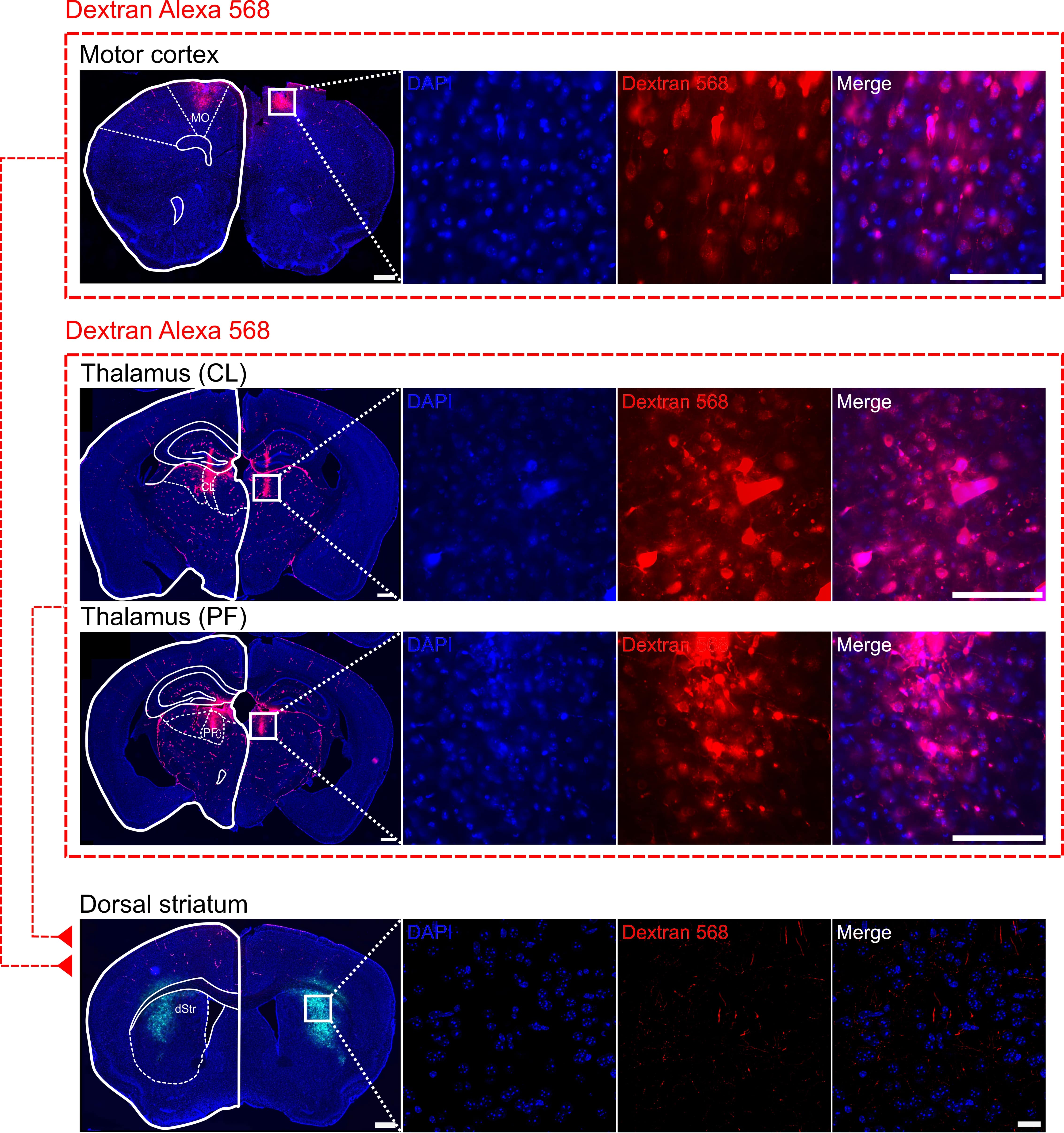

**Supplementary Fig. 3. Cortical or thalamic input-specific expression of Alexa 568-labeled dextran tracer**

Representative images of Alexa 568-labeled dextran expression in the motor cortex, thalamus (PF or CL) and dorsal striatum following cortical or thalamic input region-specific injections. The white box in each brain region (scale bars= 500 μm) is expanded views of the Alexa 568-labeled dextran expression (Motor cortex and thalamus: scale bars= 100 μm; dorsal striatum: scale bars= 10 μm).

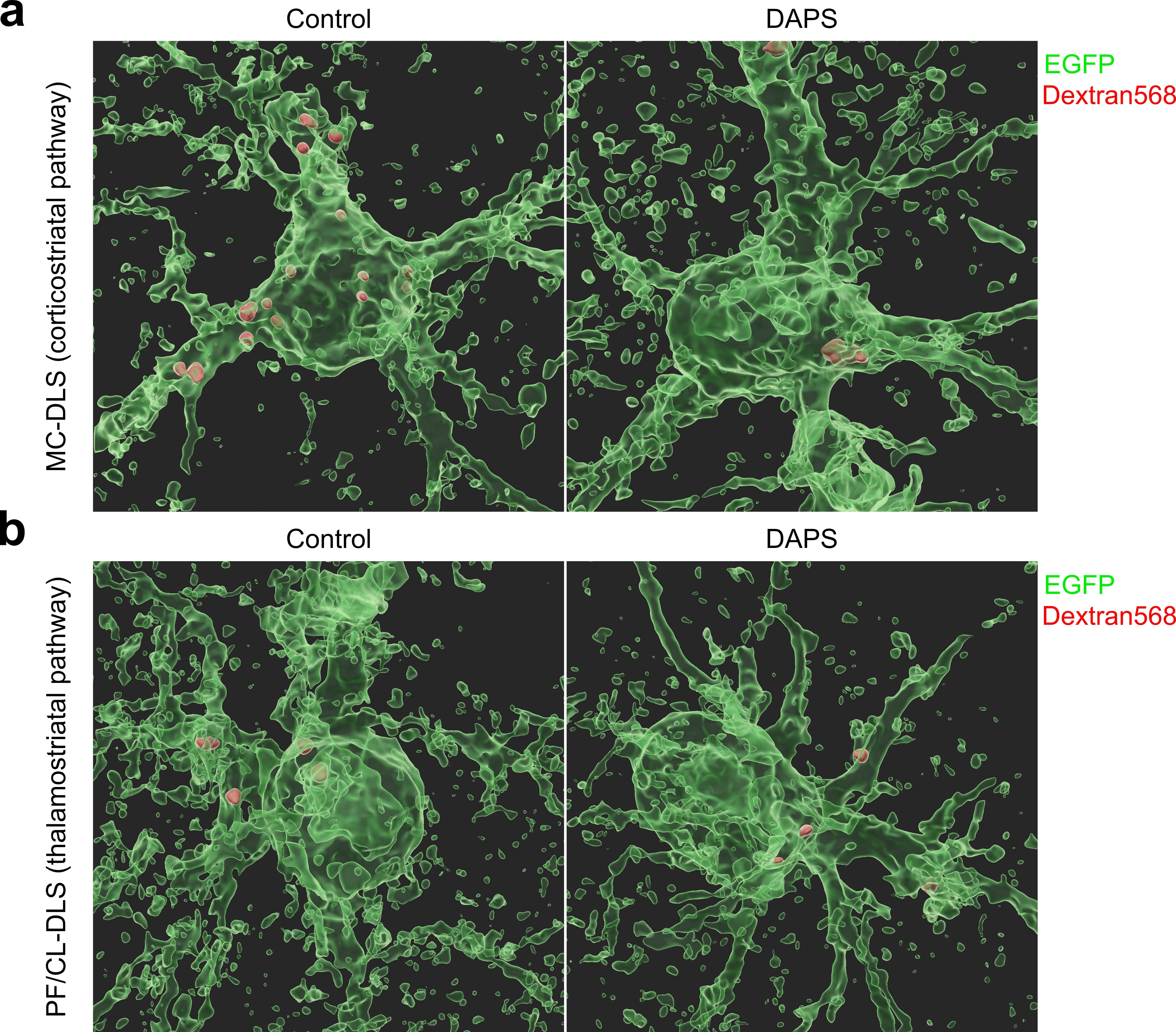

**Supplementary Fig. 4. 3D reconstructed astrocytes showing engulfment of synaptic materials**

**a-b.** Representative 3D-rendered images showing engulfed Dextran Alexa 568 puncta (red) derived from motor cortex (MC-DLS; **a**) or PF/CL (PF/CL-DLS, **b**) in astrocytes (EGFP).

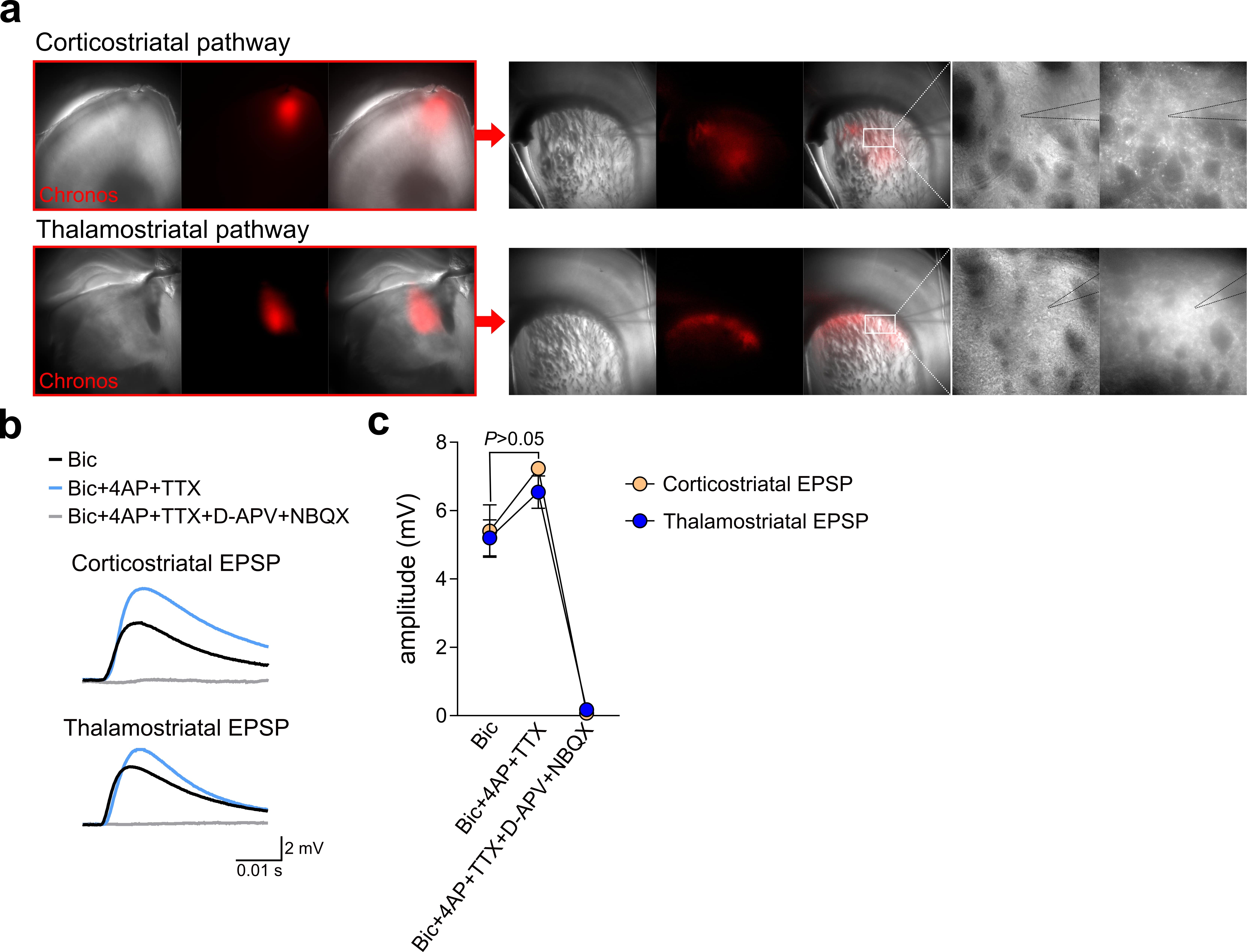

**Supplementary Fig. 5. Monosynaptic and glutamatergic responses in corticostriatal and thalamostriatal pathway-dependent optogenetic light-evoked EPSPs.**

**a.** Representative images showing corticostriatal or thalamostriatal projection-specific AAV-hSyn-Chronos-tdTomato expression and whole-cell patch clamp recording. **b.** Representative traces showing light-evoked EPSPs specific to the corticostriatal or thalamostriatal projections during drug treatment. **c.** Dot plots depicting average changes in corticostriatal or thalamostriatal EPSPs depending on each drug treatment. Corticostriatal EPSP, n= 4 cells from 2 mice; Thalamostriatal EPSP, n= 4 cells from 2 mice. Unpaired student’s t-test. Mean ± s.e.m.

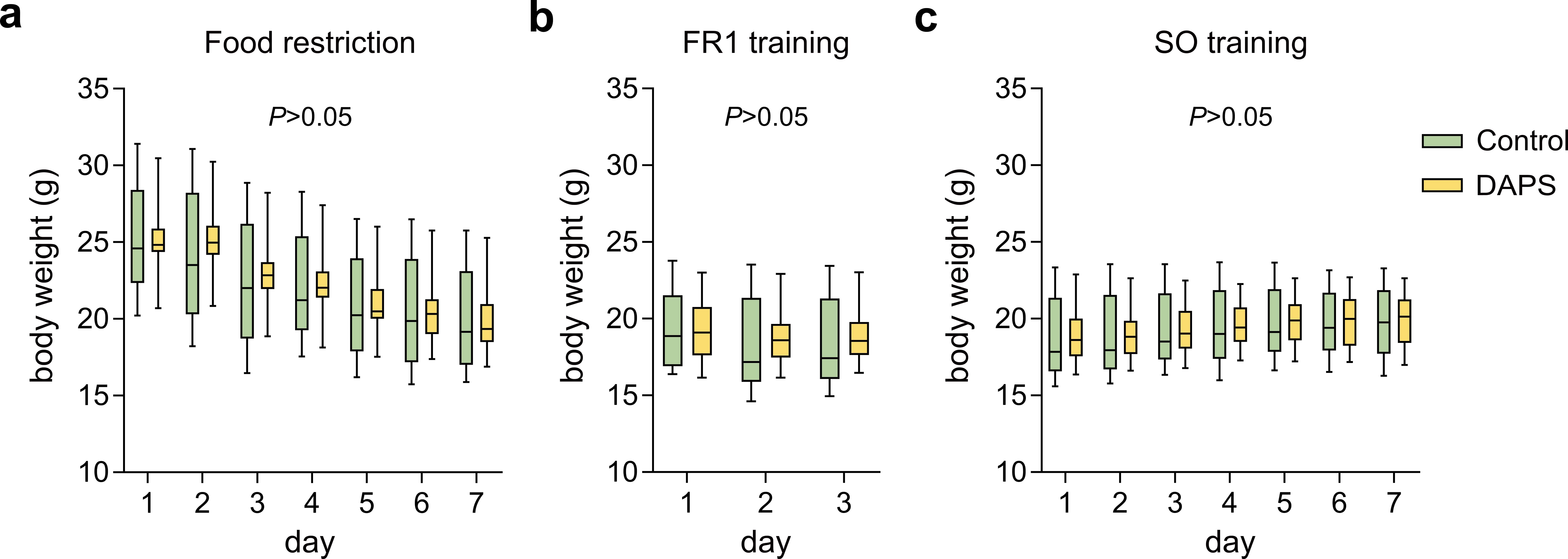

**Supplementary Fig. 6. Comparable weight loss between control and DAPS mice.**

**a-c.** Min to max plot summary data for body weight changes in control and DAPS mice during food restriction (**a**), FR1 training (**b**), and SO training (**c**). Control: n= 5 mice; DAPS: n= 10 mice. Repeated measures Two-way ANOVA test. Mean ± s.e.m.

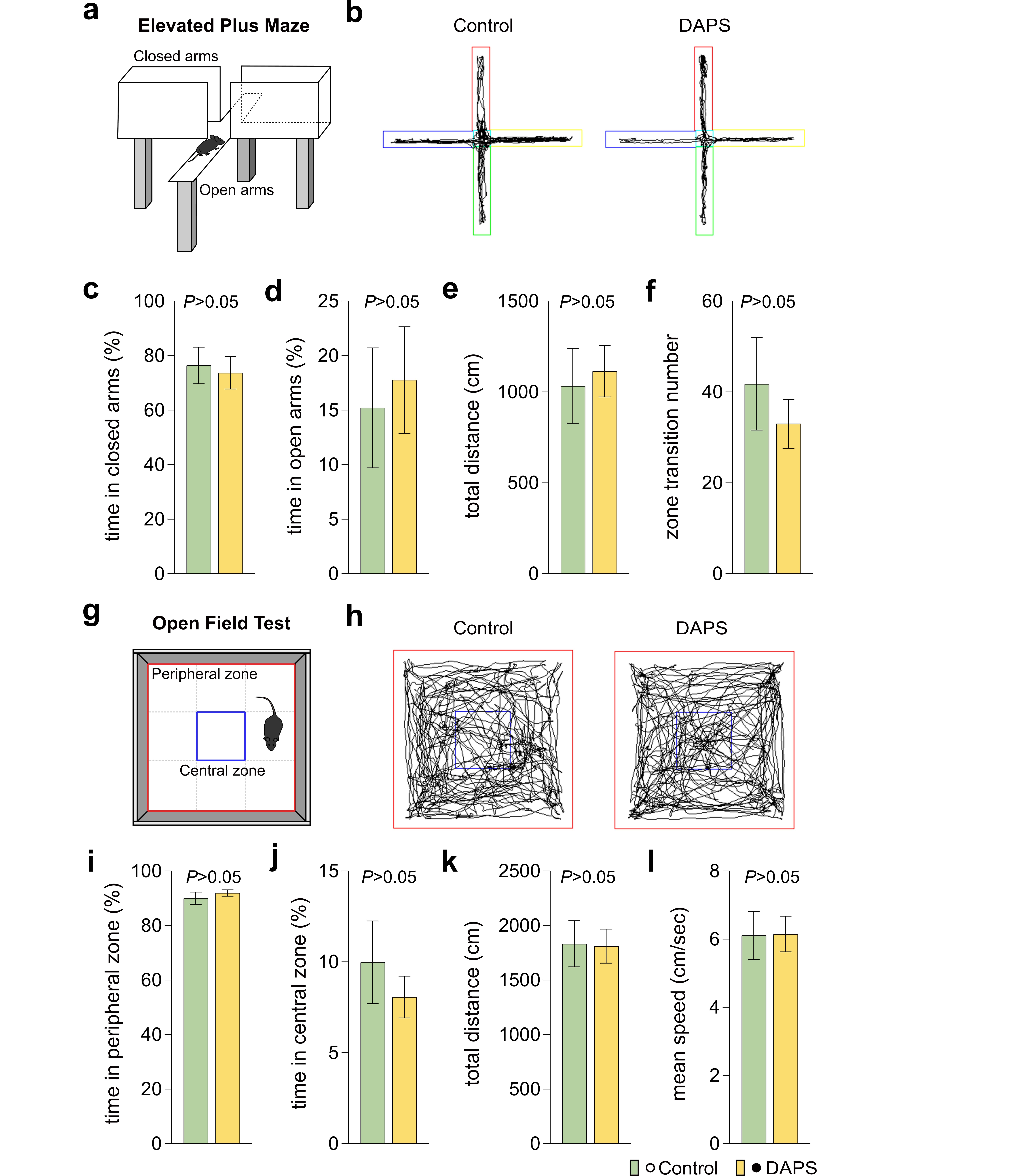

**Supplementary Fig. 7. Normal locomotion and anxiety levels in DAPS mice.**

**a.** Schematic illustration of elevated plus maze. **b.** Representative traces showing mouse-moving in an elevated plus maze in control (left) and DAPS mice (right). **c-f.** Bar graph depict percentage of time in closed arms (**c**), percentage of time in open arms (**d**), total distance (**e**), and transition number of zone (**f**) of control and DAPS mice. **g.** Schematic illustration of open field test. **h.** Representative traces showing mouse-traveling in an open field chamber in control (left) and DAPS mice (right). **i-l.** Bar graphs depict percentage of time in peripheral zone (**i**), percentage of time in central zone (**j**), total distance (**k**), and mean speed (**l**) of control and DAPS mice. Control: n= 10 mice; DAPS: n= 15 mice. Unpaired student’s t-test. Mean ± s.e.m.

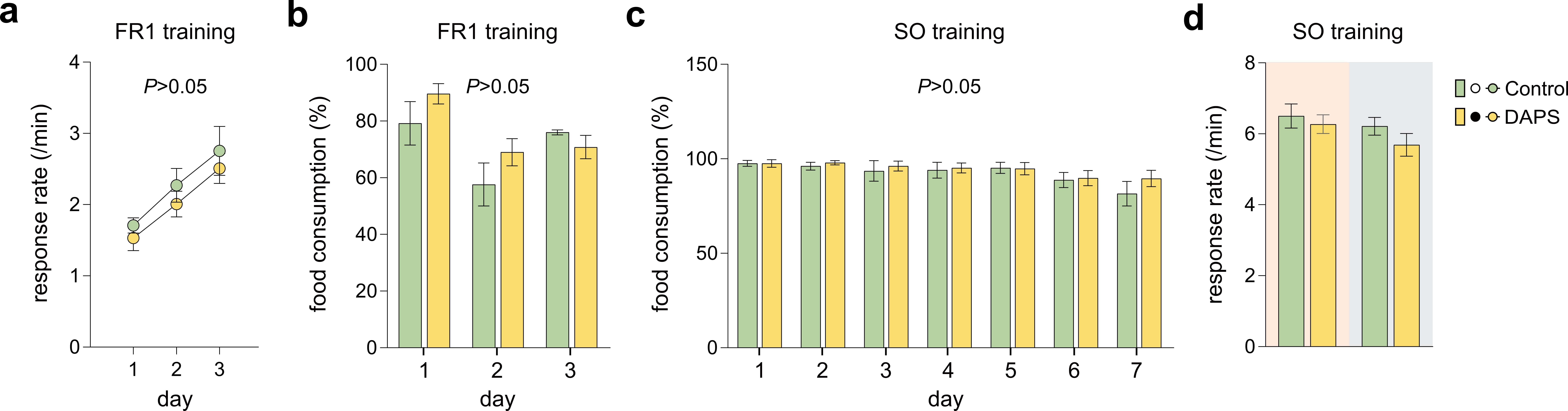

**Supplementary Fig. 8. DAPS mice have normal motivation levels and lever pressing behaviors**

**a.** Average rate of lever pressing per minute of control and DAPS mice during fixed ration 1 (FR1) training. **b-c.** Bar graphs indicate the food consumption of control and DAPS mice during FR1 training (**b**) and SO training (**c**). **d.** Bar graphs depicting average rates of lever pressing per minute during the early and late stages of SO training. Control: n= 5 mice; DAPS: n= 10 mice. Repeated measures Two-way ANOVA test. Mean ± s.e.m.

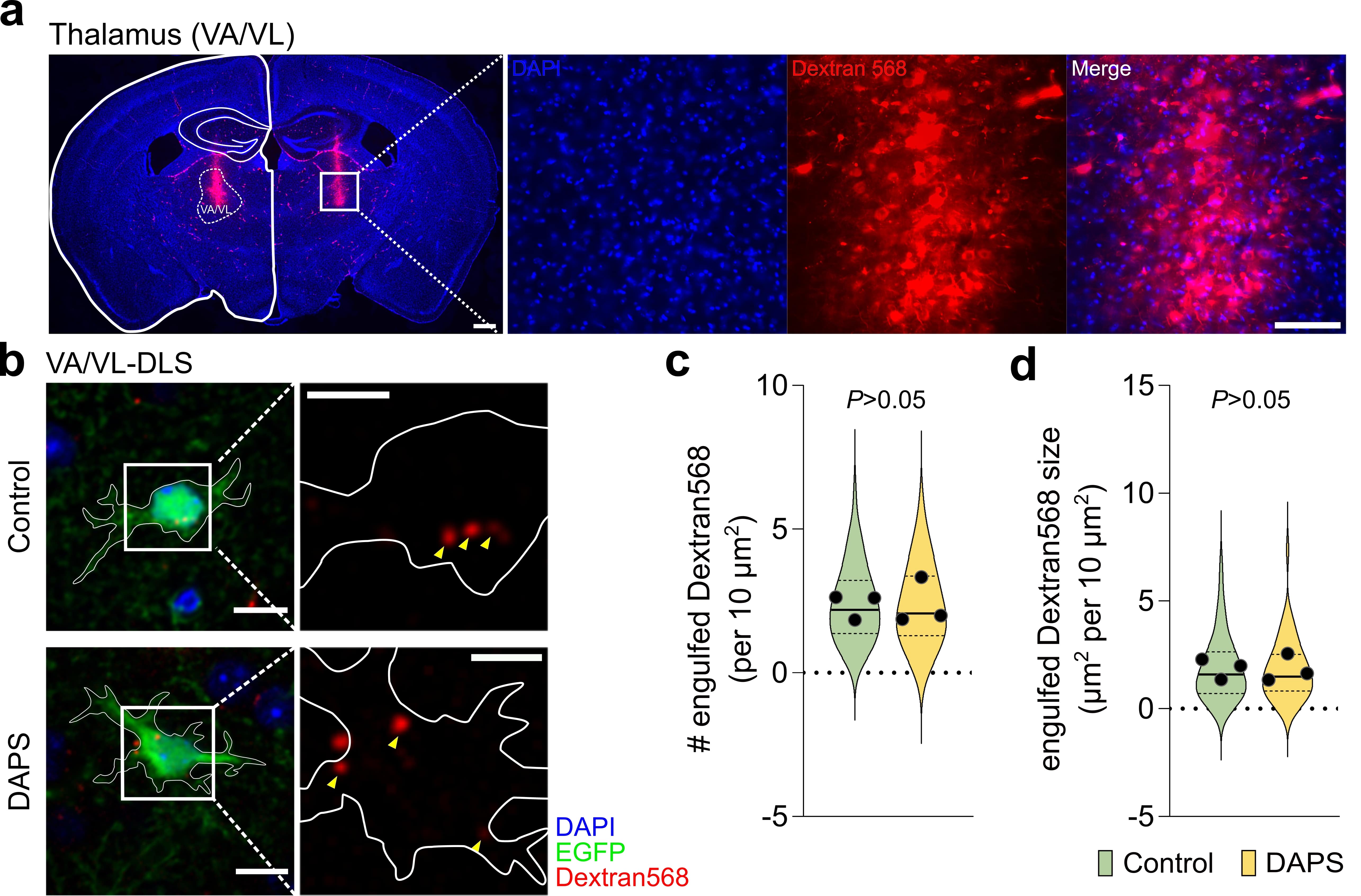

**Supplementary Fig. 9**

**Phagocytosis of VA/VL-derived thalamostriatal synapses is independent on astrocytic MEGF10/MERTK**

**a.** Representative images of Alexa 568-labeled dextran expression in the VA/VL regions. The white box in each brain region (scale bars= 500 μm) is expanded views of the Alexa 568-labeled dextran expression. Scale bars= 100 μm. **b.** Representative confocal images (left) and expanded panels (right) of astrocytes (green) with fluorescent tracers (red) derived from the ventral anterior/lateral nucleus (VA/VL) in the striatum of control and DAPS mice. The white outlines show territories of the observed astrocytes. Scale bars= 10 μm. Scale bars= 5 μm in the expanded panels. **c-d.** Summary of the number (**c**) and size (**d**) of phagocytosed fluorescent tracers found in the astrocytic cytosol in control and DAPS mice. Control, n= 45 images from 3 mice; DAPS, n= 44 images from 3 mice. Unpaired student’s t-test. Mean ± s.e.m.

**Table 1. Electrophysiological properties of the recorded MSNs**

|  | **Membrane capacitance**  **(pF)** | **Membrane resistance (mΩ)** | **Resting membrane**  **(mV)** |
| --- | --- | --- | --- |
| **Control** | 35.06 ± 3.72 | 156.70 ± 9.77 | -74.00 ± 1.20 |
| **DAPS** | 43.70 ± 4.75 | 140.10 ± 7.95 | -75.30 ± 1.43 |
| **P value** | *P*= 0.1664 | *P*= 0.1933 | *P*= 0.4948 |
